# Deformity Index: A semi-reference quality metric of phylogenetic trees based on their clades

**DOI:** 10.1101/706440

**Authors:** Aritra Mahapatra, Jayanta Mukherjee

## Abstract

Measuring the correctness of a phylogenetic tree is one of the most fundamental tasks in phylogenetic study. A large number of methods have been proposed to measure the correctness of a tree. Such methods completely depend on the reference tree and they compute the distance between reference the tree and the target tree. But it is very difficult to obtain a precise and an accurate reference tree for a selected dataset. As a result, the existing methods for comparing the phylogenetic trees can behave unexpectedly in various scenarios. In this paper, we introduce a scoring function, called the Deformity Index, to measure the correctness of a tree based on the biological knowledge of the clades. The strength of our proposed method is that it does not consider any reference tree. We have also investigated the range and the distributions of the different modules of Deformity Index. Furthermore, we perform different goodness of fit tests to understand its cumulative distribution. We have also examined in detail the robustness as well as the scalability of our measure by different statistical tests under the Yule and the uniform models. Moreover, we show that our proposed scoring function can overcome the limitations of the conventional methods of tree comparing by experimenting on different biological datasets.

## 1 Introduction

A phylogenetic tree represents the traits of evolution between a set of species [12]. The leaf nodes are the modern day species of known characteristics, whereas the internal nodes are considered as the hypothetical ancestors.

In studying phylogeny of different species, both of the phenotype characters (such as morphologcial features, living habits, etc.) as well as the genotype characters (such as genome sequences, gene sequences, exons, introns, etc.) are being considered [45, 7, 20, 5, 6, 49, 24]. There are various methods for constructing phylogenetic trees from the genotype data. These methods are broadly classified into two groups, i.e., alignment based methods and alignment free methods. The alignment based methods are further classified into four sub-categories, i.e., distance based, parsimony based, maximum likelihood based, and Bayesian methods [12, 21]. Alignment free methods are categorized into four subgroups, such as, k-mer frequency based [46], substring based [54], information theory based [15], and graphical representation based methods [1].

Modern advent in genotyping and sequencing techniques is capable of producing different genetic sequences with a decreasing cost [13, 35] but with an advanced phylogeny research. However, different types of data and methods propose different phylogenetic relationships among the species. For example, more than hundred studies are underway on Gadiformes since 1903 [39] which have proposed different phylogenetic relationships [10, 41, 45]. In this situation, different methods have been proposed to compute the correctness of the derived phylogenetic trees with respect to a reference tree. Some of the popular methods are Robinson-Fould (RF) distance [42], Matching Split (MS) distance [2], Nodal Split (NS) distance [3], Triples (TT) distance [8], etc.

Robinson-Fould distance [42] computes the distance between two trees by considering all of the bipartitions in each of them. Let Φ(*T*) denote the set of all of the bipartitions of a tree *T*. The Robinson-Fould distance between two trees *T*_1_ and *T*_2_, say *d_RF_* (*T*_1_, *T*_2_), is the half of the cardinality of the symmetric difference between Φ(*T*_1_) and Φ(*T*_2_), hence,

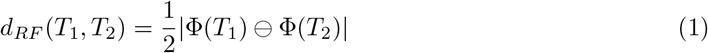

where, Φ(*T*_1_) and Φ(*T*_2_) are the bipartitions of *T*_1_ and *T*_2_, respectively and |*χ*| is the cardinality of the set *χ*.

Similar to the RF distance, Matching Split distance [2] considers the bipartitions (also called splits) of the trees, say (*A*|*B*). For trees *T*_1_ and *T*_2_, let, the set of bipartitions be denoted as Φ(*T*_1_) = S(*A*_1_|*B*1) and Φ(*T*_2_) = *S*(*A*_2_|*B*_2_), respectively. Considering the bipartitions (*A*_1_|*B*1) and (*A*_2_|*B*_2_) as two disjoint sets, a complete bipartite graph (*K*_2_,_2_) can be formed. Hence, the set of all of the complete bipartite graphs, *S*(*K*_2_,_2_) = S(*A*_1_|*B*1) × *S*(*A*_2_|*B*_2_). For such a complete bipartite graph *k*, *k* ∈ *S*(*K*_2_,_2_), with two disjoint sets 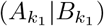 and 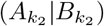, the minimum-weight perfect matching, *ω*(*k*), can be represented as,

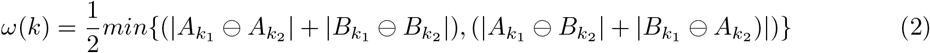

Let for Φ(*T*_1_) and Φ(*T*_2_) the minimum-weight perfect matching is *M*. The Matching Split distance between two trees *T*_1_ and *T*_2_, say *d_MS_* (*T*_1_, *T*_2_), is the total weight of a minimum-weight perfect matching (*M*),

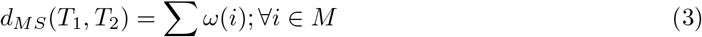

The Nodal Split (NS) distance [3] uses the concept of the split path lengths matrices of a tree. The path between two leaves *i* and *j* of a tree *T* is split by their most recent common ancestor (say *ij*). The split path lengths of two leaves *i* and *j* of a tree *T*, denoted as *P*_*T*_ (*i*, *ij*) and *P*_*T*_ (*j*, *ij*), respectively, are the length of the paths from the most recent common ancestor of *i* and *j* to node *i* and node *j*, respectively. Considering all of the the pairwise nodes of the tree, the *P*_*T*_ form the matrix called split path lengths matrix. The split path lengths matrix is denoted as 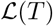. The NS distance between the two trees *T*_1_ and *T*_2_ is represented as,

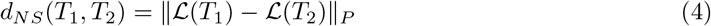

where, ||*χ*||*P* is the *L*^*P*^ norms of the matrix *χ*. In general, NS distance considers either Manhattan norm (*p* = 1) or Euclidean norm (*p* = 2) to compute the distance between the two trees.

The Triples (TT) distance [8] counts the number of the triplets which are different in the two trees.

## 2 Limitations of the conventional tree comparison methods

### 2.1 Unknown relations - Multifurcation

The existing conventional tree comparison methods are completely dependent on the reference trees. But a complete knowledge of the reference tree for the selected species of a study may not be available. It is also very difficult to get the knowledge of some evolutionary relationships among the selected species. In such cases, we assume multifurcation at the MRCA of the selected species (Fig. 1(a)). Existing tree comparison methods are very strict towards the relationships among the species. Hence, it shows its limitation when multifurcation occurs. A typical example is illustrated in Fig. 1. Let us consider the tree in Fig. 1(a) as the reference tree which have a multifurcation at a node (refer to the blue colored clade in Fig. 1(a)). The same set of species forms monophyletic clade in the two trees shown in Fig 1(b, c). Though both of the trees are having the same monophyletic clade as that in the reference tree, still the conventional tree comparing methods report non-zero scores due to the presence of false positive clades (the clades which are present in the target trees but not in the reference tree) (Fig. 1(a)). In this view, it may be observed that counting only the false negative clades (those which are present in the reference tree but not in target trees) provides better result. On the contrary counting only the false negative rate also has its limitation which we describe in the next section.

**Figure 1:**
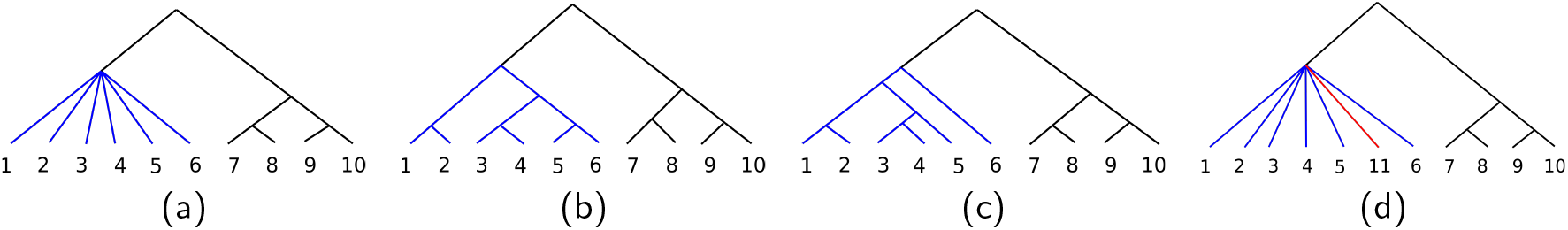
An example representing the limitations of the conventional tree comparing methods. (a) Reference tree having multifurcation at the node represented by blue colored clade. (b) The tree represents a clade, shown in blue color, having the same set of species as that of in the multifurc clade of the reference tree. But the speciations of the species are known here. The *RF* = 2, *MS* = 10, *NS* = 10.95, and *TT* = 20 w.r.t. the reference tree. (c) The *RF* = 2, *MS* = 12, *NS* = 13.27, and *TT* = 20 w.r.t. the reference tree, (d) The tree having an outgroup species (11), shown in red color, within the blue clade. The species 11 is not present in the reference tree. The *RF* = *MS* = *NS* = *TT* = 0 w.r.t. the reference tree as the additional taxa (11) has been pruned.

### 2.2 Unavailability of the absolute reference tree

It is to be mentioned that the extension of the phylogenetic tree is very much necessary for the success of the *Tree of Life Web Project* [16]. But all of the tree comparing measures work only when both of the reference tree and the target trees have the same set of species. Let us consider a case where the reference tree and the target tree have different sets of species. Let Fig. 1(a and d) be the reference and the target trees, respectively. Consider that the target tree (refer to Fig. 1(d)) has an outgroup species (11) (shown in red color) within the blue clade while this species (11) is not present in the reference tree (please refer to Fig. 1(a)). So by observing it can be said that the target tree topology violates the reference topology. But the conventional methods first prune the species 11 to ensure that both of the trees have the same set of species which also makes the trees topologically identical. Hence, all the measures reflect zero scores though the blue clade of Fig. 1(d) violates its monophyletic nature based on the reference clade. Counting the false positive rate in this case gives more meaningful result than that of the false negative rate. Hence, any one of the false postive rate and false negative rate as a single measure is not sufficient, and it is inconvenient to compare two techniques by the combination of false positive rate and false negative rate.

Due to these limitations of the conventional measures, most of the recent phylogenetic studies measured the correctness of their proposed tree by using the following methods,

1. ***Manual inspection***. The correctness of the derived trees is checked manually by validating the presence of each clade using biological knowledge [25]. But this technique does not provide any quantitative measure.
2. ***Comparing with other methods***. In this approach, a phylogenetic tree is formed by applying other established methods on the same set of species and consider the tree as the reference tree for the corresponding dataset [61]. The problem of this approach is that, we have to consider the established methods as the gold standard which are not beyond contentions.
3. ***Indirect method***. This approach computes different errors, like root mean square error of distance matrices [58, 62]. So this method overlooks the accuracy of the final tree construction steps. There are some methods like maximum likelihood, Bayesian estimation, which can directly derive trees from the sequences without generating distance matrices. For this case the indirect method will not be applicable.

In this regard, we propose a method to provide a quantitative measure of the correctness of a phylogentic tree based on the biological knowledge of the different clades of the tree. We call it as ***Deformity Index***. Finally, the cumulative score of the clades gives the Deformity Index of the tree. The computation of Deformity Index only depends on the given clades, so the relationships within a clade do not affect the computation of Deformity Index. Hence, this method does not consider the precise relationships among the species within a clade. So it can be applicable for all types of trees irrespective of bifurcation or multifurcation. We believe that, this is the first method to measure the correctness of a tree without using a reference tree.

## 3 Basic Facts and Definitions

Let *G* = (*V*, *E*) is a *graph* with a set of vertices *V* and a set of edges *E*. An acyclic connected graph is considered as the *tree*. *Phylogenetic tree* depicts the evolutionary relationships among a set of species *S*. If the set of leaves of the phylogenetic tree is *L*, then *L* → *S* is a bijective function [12].

### Definition 1. Rooted phylogenetic tree. [12]

*A phylogenetic tree in which a nonleaf vertex is considered as the **root** of the tree.*

In a phylogenetic tree, *T*, the nonleaf vertices, *V*_*I*_ = *V* \ *L*, are unlabeled and are considered as the *hypothetical ancestors*. The minimum degree of the root node (**r_T_**) is two, whereas, other *V*_*I*^s^_ have degrees greater than or equal to three. If the degree of these vertices is strictly three then this tree is called a *rooted binary phylogenetic tree* or a *resolved phylogenetic tree*. Whereas, a vertex degree greater than three (or having more than two descendants) denotes the *multifurcation* at that vertex. A tree having multifurcation is considered to be an *unresolved phylogenetic tree* which reflects that the exact relationships among those species are not clear. In Fig. 2 the speciation point denoted by orange dot represents the multifurcation. A rooted binary phylogentic tree that reduces to a path after deleting all the pendent edges and the leaves is called a *caterpillar tree*.

The list of all leaves under an internal node *η*, ∀*η* ∈ *V_I_*, of a tree is represented as *λ*(*η*). Except the root node, **r_T_**, the parent node or the recent ancestor of a node *n*, ∀*n* ∈ *V* \ *r_T_*, is represented as *p*(*n*). For an example, let us consider the internal nodes *η_V_* and *η_O_* which are represented as violet and orange dots, respectively, in Fig. 2. Then, *λ*(*η_V_*) = {*A, B, C, D, E*} and *λ*(*η_O_*) = {*M, N, O*}. For the leaf node *N*, the parent node, *p*(*N*) is represented as orange dot, in Fig. 2.

**Figure 2:**
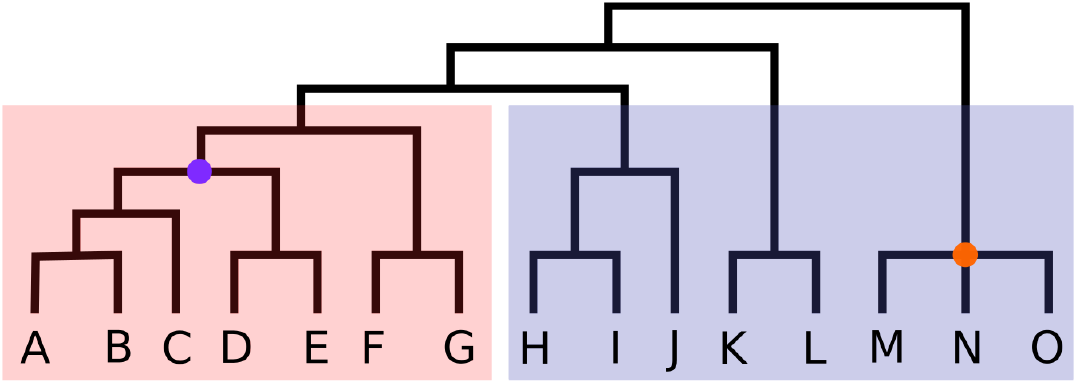
Rooted phylogenetic tree. The violet dot is the most recent common ancestor of the species *A*, *B*, and *D*. All of the descendants of the green node, *A*, *B*, *C*, *D*, and *E*, form a clade. Let us consider that the tree has two taxonomic groups of species denoted as red and blue color. The red group of species forms a monophyletic clade, whereas the blue group species forms a paraphyletic clade.

### Definition 2. Level

*The number of edges between the root node and a node is considered as the level of the corresponding node*.

The level of a node *n* is represented as *h*(*n*). The level of the root of a tree is considered as one. In Fig. 2, the level of the violet node (*η_V_*) and orange node (*η_O_*) are *h*(*η_V_*) = 5 and *h*(*η_O_*) = 2, respectively.

### Definition 3. Height

*The number of edges between a node and its furthest descendant leaf node is considered as the height of the corresponding node*.

The height of a node *n* of a tree is represented as *H*(*n*). The height of the leaf nodes are considered as zero. The height of the root of the tree is considered as the height of the tree. In Fig. 2, the height of the violet node (*η_V_*) and orange node (*η_O_*) are *H*(*η_V_*) = 3 and *H*(*η_O_*) = 1, respectively. Whereas the height of the tree in Fig. 2 is 7.

### Definition 4. Most recent common ancestor. [12]

*For a set of species from a rooted phylogenetic tree, the most recent internal node from which all those species of the set are descended is called the most recent common ancestor*.

Most recent common ancestor (MRCA) of a group is the speciation point of the group. In Fig. 2, for the list of species, t = {*A, B, C, D, E*}, t ∈ *L*, the MRCA of 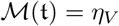, represented as violet dot in Fig. 2.

### Definition 5. Taxonomic group. [4]

*Organisms are grouped on the basis of their common characteristics. These groups are given a taxonomic rank. Groups of taxonomic ranks assemble and form the higher rank or a super-group, thus forming a taxonomic hierarchy*.

There are nine principal ranks of taxonomy, i.e., life, domain, kingdom, phylum, class, order, family, genus, and species. The domain is the highest rank and species is the lowest rank among the principal taxonomic ranks. It is to be mentioned that a group of organisms from the higher rank share more general characteristics while the group of organisms from the lower rank share more specific characteristics. For an example, dog, cow, and dolphin both belong to the same kingdom (Animalia), phylum (Cordata), and class (Mammalia), whereas their other lower taxonomic ranks are different.

### Definition 6. Clade. [12]

*The set defined by all of the descendant leaves of an internal node is called as clade*.

A clade is a subtree of the original tree. A clade is represented as the list of all leaves present under the node. Hence, *λ*(*η*), ∀*η* ∈ *V*_*I*_, is a clade. So, it is to be mentioned that, 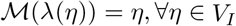

It is assumed that all of the members of a clade share some common and distinct traits. If all the members of a clade belong to the same taxonomic group then that clade is called a ***monophyletic clade***, otherwise, the clade is called a ***paraphyletic clade*** (Fig. 2).

## 4 Methodology

The proposed measure computes the correctness of a tree by considering the information of the reference clade(s). A phylogenetic tree can be represented as a set of clade(s) where each clade represents a list of leaves under an internal node (please refer to Fig. 3). Hence in our method, providing a reference tree is also equivalent to provide a list of clades of that reference tree.

**Figure 3:**
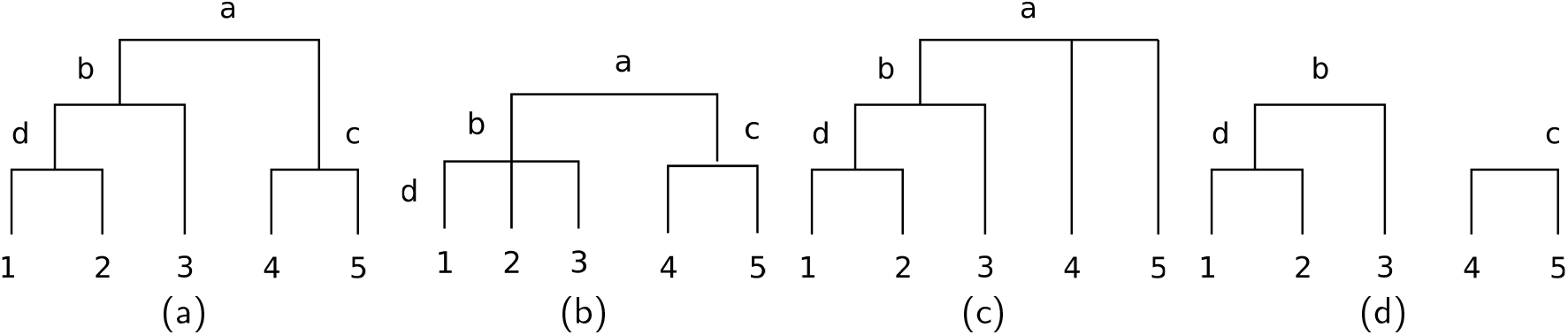
An example which represents the clades of the tree in different cases. (a) The tree is a binary tree and have complete information about all of its clades. Λ(*T*) = {{1, 2, 3, 4, 5}, {1, 2, 3}, {1, 2}, {4, 5}}. (b) The more precise information about the 1, 2, and 3 within the clade *b* is not present, hence the node *b* become multifurced. The Λ(*T*) = {{1, 2, 3, 4, 5}, {1, 2, 3}, {4, 5}}. (c) Here, Λ(*T*) = {{1, 2, 3, 4, 5}, {1, 2, 3}, {1, 2}}. The precise information about the relation between 4 and 5 is not known, hence the node *a* become multifurced. (d) Here, Λ(*T*) = {{1, 2, 3}, {1, 2}, {4, 5}}. As information of the clade *a* is not there, so *T* is being considered as two components.

Our objective is to compute degree of deformation of the clades in a phylogenetic tree with respect to the reference clades.

### Definition 7. Transfer In

*The distance (number of edges) required to shift a species to attach it within a selected clade is called Transfer In*.

The movement from one node to another is applicable only for the leaves of the tree. The *Transfer In* of a species is required when we need to join a species to its correct node. If we add a species *A* at the node *η*, then the species *A* is required to move from *p*(*A*) to *η*. The **Transfer In** for moving species *A* to *η* is denoted as *T L_in_*(*A*, *η*). Hence,

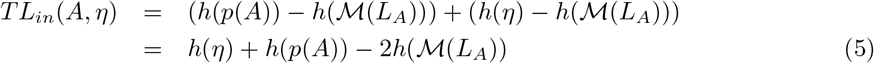

where *L*_*A*_ = *λ*(*η*) ∪ *A*; ∀*A* ∈ *L* and ∀*η* ∈ *V*_*i*_.

For an example in Fig. 4(a), to add the species 4 of the tree, *T*, at the mentioned clade (*η*), it needs to move two edges. So, *T*_*Lin*_(4, *η*) = 2.

**Figure 4:**
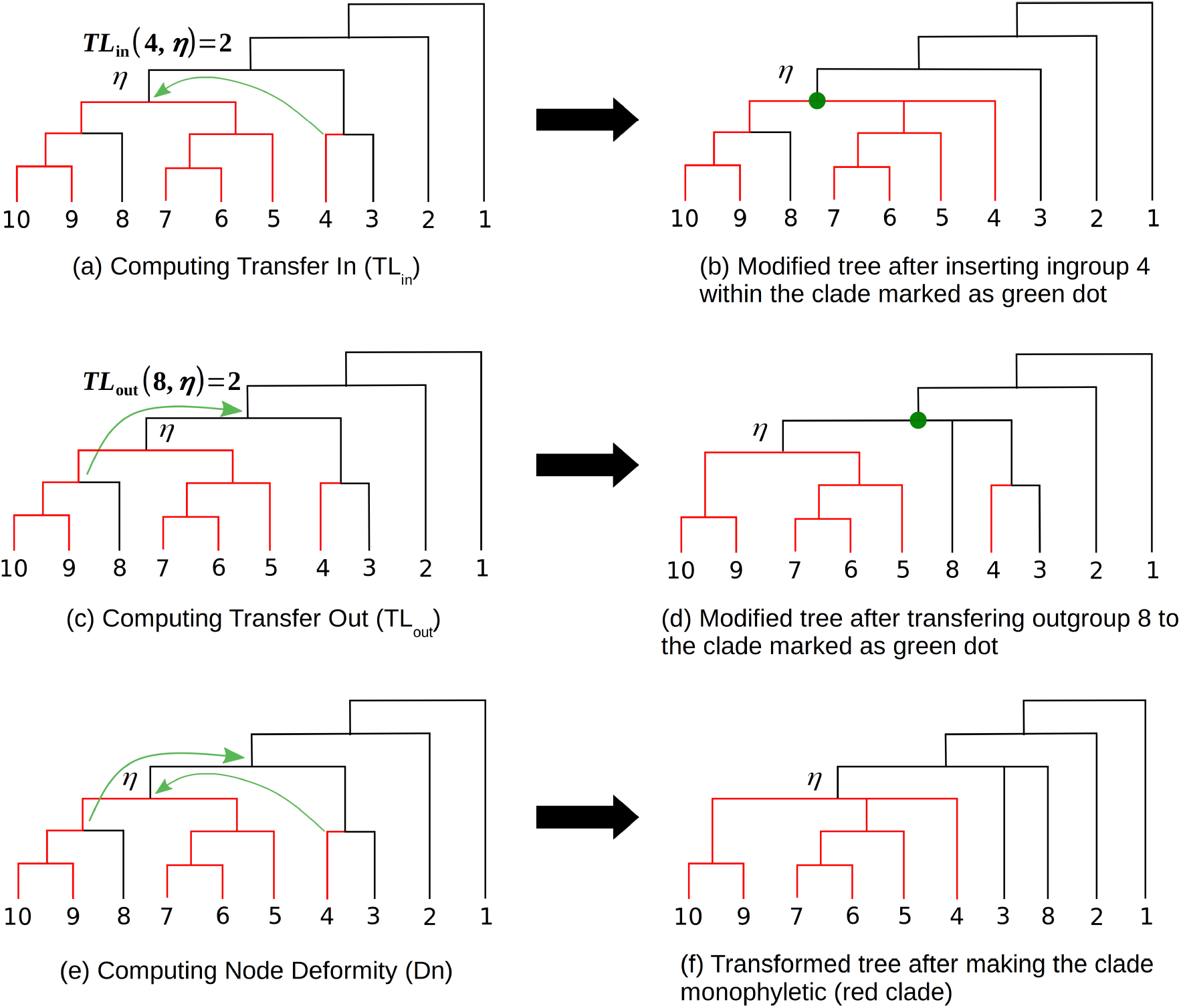
Consider a reference clade, Λ_*i*_ = {4, 5, 6, 7, 9, 10} whose members are denoted as red color. (a) **Transfer In:**To make the clade *η* monophyletic with respect to the Λ_*i*_, species *4* should be placed within the clade *η*. Hence the species *4* should be moved two labels (marked as green arrow). So the *TL*_*in*_(4, *η*) = 2. (b) The modified tree after adding the ingroup species *4* to the clade *η*. (c) **Transfer Out:**To make the clade *η* monophyletic with respect to the Λ_*i*_, species *8* should be placed outside of the clade. It requires to be shifted two levels from the current position (marked as green arrow). So the *TL*_*out*_(8, *η*) = 2. (d) The modified tree after removing the outgroup species *8* from the clade *η*. (e) The **Node Deformity**is the sum of all Transfer In and Transfer Out. Hence, the *Dn*(*η,* Λ_*i*_) = (2 + 2) = 4. The **deformation**, 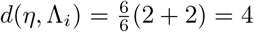. (f) Transformed tree after applying *Dn*(*η,* Λ_*i*_).

### Definition 8. Transfer Out

*The distance (number of edges) required to shift a species to remove it from a selected clade is called the Transfer Out*.

Similar to the Transfer In, only the leaf node can be removed from a particular clade. The *Transfer Out* of a species is required when a species is required to be removed from a clade. If we remove a species *B* from a clade *η*, then the species *B* is required to be moved from *p*(*B*) to outside of the *η*. The **Transfer Out**of the species *B* is denoted as *TL*_*out*_(*B*, *η*). Hence,

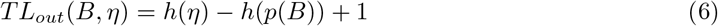

where, ∀*B* ∈ *L* and ∀*η* ∈ *V_I_*

For example, in Fig. 4(c), to remove the species 8 of the tree, *T*, from the mentioned node (*η*), it needs to move two edges. So, the Transfer Out, *TL*_*out*_(8, *η*) = 2.

Consider a tree, *T*, having the leaves *L* and the internal nodes *V*_*I*_. To compute the Deformity Index of the target tree, *T*, we have a list of reference clades, say Λ(*TR*), as the input of this process. Consider a reference clade, Λ_*i*_: Λ_*i*_ ∈ Λ(*T*_*R*_). At any particular node of the target tree, *η* ∈ *V*_*I*_, it may be required to add a list of species, *P*, where, *P* = Λ_*i*_ \ *λ*(*η*) and to remove a list of species, *Q*, where, *Q* = *λ*(*η*) \ Λ_*i*_ to make the clade of the target tree monophyletic at the internal node *η*. For each case, we can compute the Transfer In for each *p* ∈ *P* and the sum of the Transfer Out for each *q* ∈ *Q*, respectively. We consider the total of them and call them as *Total Transfer In* and *Total Transfer Out*, respectively. At an internal node *η*, the *Total Transfer In* and *Total Transfer Out* of the reference clade Λ_*i*_ are represented as *TTL*_*in*_(*η,* Λ_*i*_) and *TTL*_*out*_(*η,* Λ_*i*_), respectively. Hence,

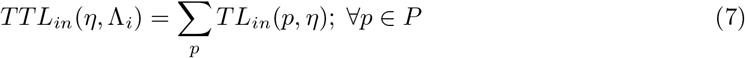

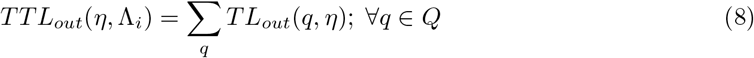

For the reference clade Λ_*i*_, the sum of *TTL*_*in*_(*η,* Λ_*i*_) and *TTL*_*out*_(*η,* Λ_*i*_) is considered as the ***Node Deformity*** of Λ_*i*_ at *η*. This *Node Deformity* is denoted as *Dn*(*η,* Λ_*i*_). The *Dn*(*η,* Λ_*i*_) is normalized by the fraction of the size of the reference clade (Λ_*i*_) and is considered as ***deformation*** of Λ_*i*_ at *η*. This *deformation* is denoted as *d*(*η,* Λ_*i*_). Hence,

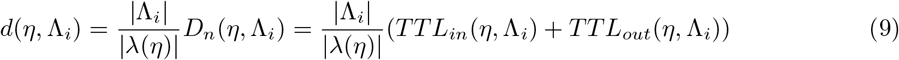

where, |**χ**| symbol represents the number of members present in the list **χ**.

Now, our objective is to compute the minimum deformation of each reference clade, Λ_*i*_: ∀Λ_*i*_ ∈ Λ(*T*), within the target tree, *T*.

### Definition 9. Clade Deformation

*The Clade Deformation of a reference clade within the target tree is the minimum deformation of the clade among all internal nodes*.

The Transfer In and Transfer Out make the target clade monophyletic. The minimum cost required for Total Transfer In and Total Transfer Out is the *Clade Deformation* of the target tree corresponding to the reference clade. The *Clade Deformation* with respect to Λ_*i*_ is denoted as, *Dc*(Λ_*i*_). Hence,

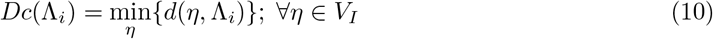

### Definition 10. d-node

*The node where the minimum of the deformation occurs for a reference clade is called here as d-node*.

It is a function of clade Λ_*i*_ and is denoted as **d**_*η*_ (**Λ**_**i**_): Λ_*i*_ ∈ Λ(*TR*). We use the term d-node in the rest of the paper to indicate the node where the Clade Deformation occurs.

For each Λ_*i*_ ∈ Λ(*TR*), we compute the *Dc*(Λ_*i*_). The ***Deformity Index*** of a tree, *T*, is the average of the Clade Deformations for all the reference clades. The *Deformity Index* of the tree, *T*, is denoted as *D*(*T*). Hence,

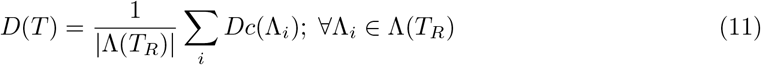

The procedure of computing deformity index is described in Fig 5.

**Figure 5:**
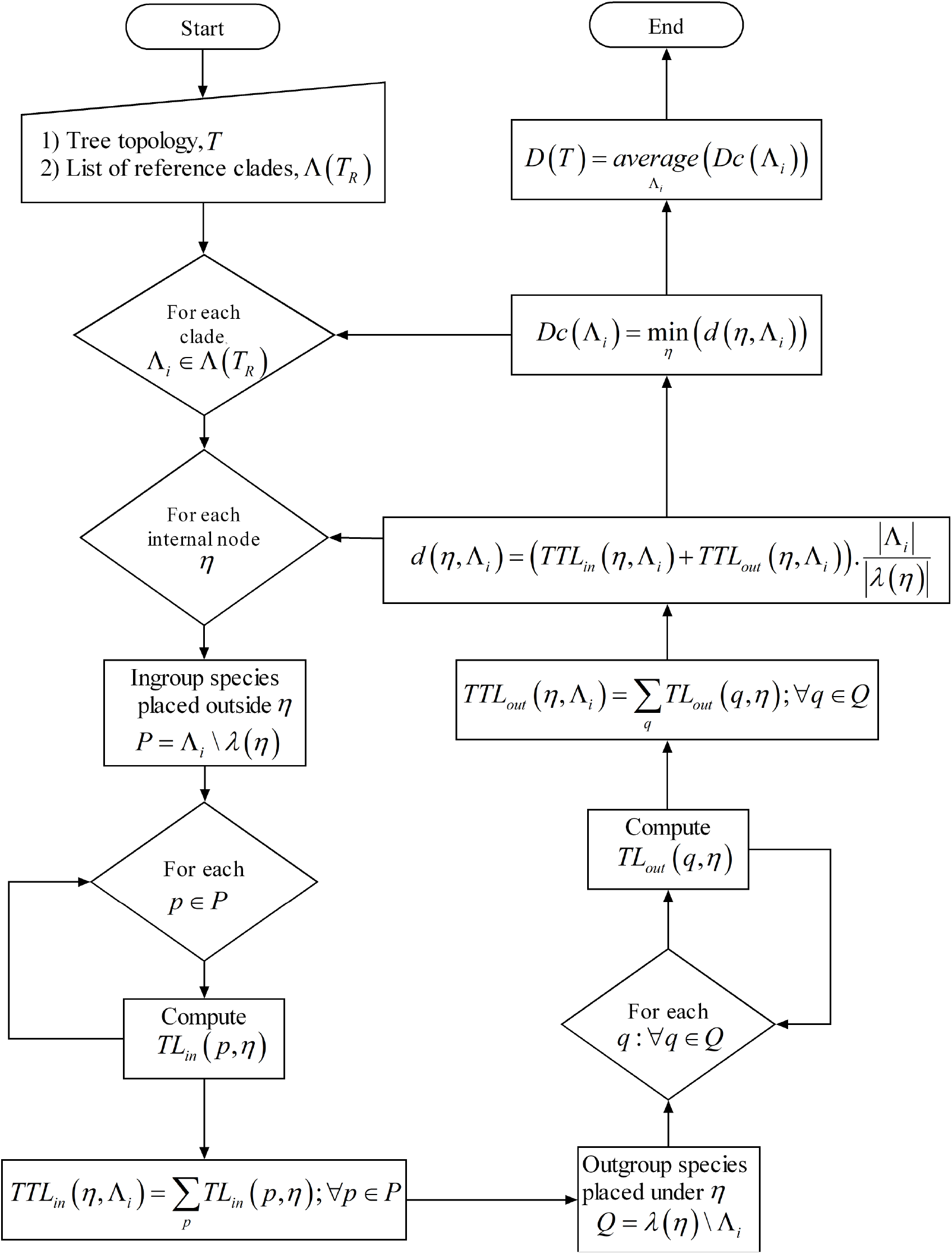
Flow diagram for computing Deformity Index of a phylogenetic tree

### 4.1 Properties of Deformity Index

In this section we derive different properties of Deformity Index, especially the maximum possible value of the Deformity Index for a given number of leaves. It is to be noted that the minimum possible value of Deformity Index is zero and it occurs when all the clades of the target tree satisfy the reference clades.

As the Deformity Index for a given reference clade depends on the topology of the target tree, hence our first objective is to identify the topology for which the value of the Deformity Index can reach its maximum level.

#### Lemma 1.

*The maximum value of the level of a node can occur if and only if the ancestors of the node of the tree are bifurcated*.

#### Lemma 2.

*For a given number of leaf nodes, the highest value of the sum of the levels of all the leaves of a tree is achieved when the tree is a caterpillar tree*.

#### Lemma 3.

*For a caterpillar tree, the sum of the levels of the elements of reference clade is inversely proportional to the clade distortion within the tree*.

For example let a caterpillar tree of *n* leaves. Considering a reference clade of *m* members. The caterpillar tree agrees the reference clade if the members form a clade in the caterpillar tree. From the caterpillar tree topology it can be understood that the members of the reference clade required to be attached at their highest possible level in the caterpillar tree to form a clade. Any distortion in the caterpillar tree reduces the level of the corresponding member.

The proof of Lemma 6, 7, and 8 are given in Appendix. From Lemma 6, 7, and 8 it can be understood that the value of Deformity Index will be maximum when the target tree is a resolved caterpillar tree and the elements of the reference clade are placed at their lowest possible level (that means nearest to the root node). Lemma 9 defines a property of the selected tree topology which is required to compute the level of d-node for a given reference clade.

#### Lemma 4.

*For a given reference clade, the deformation at each level of a caterpillar tree is a convex function*.

The proof of Lemma 9 is provided in Appendix.

#### Lemma 5.

*For a caterpillar tree of *n* number of leaves and a reference clade of size *c* (1 < *c* < *n*), the level of the d-node is *p*, where*,

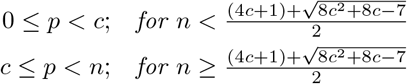

The proof of Lemma 10 is given in Appendix. Finally, in Theorem 1 we use Lemma 10 to compute the level of d-node for a given reference clade.

#### Theorem 1.

*Among the trees of *n* leaves, the level of the d-node where the maximum of the Clade Deformation occurs for the reference clade of c* (1 < *c* < *n*) *number of elements is denoted as h*(*dη* (Λ_*R*_)), *then*,

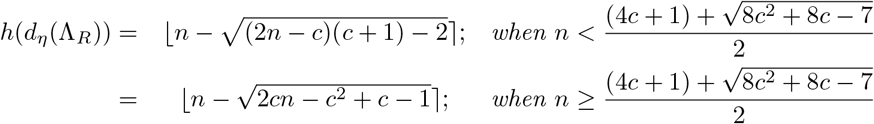

*where*, 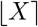 *denotes the nearest integer value of X*.

#### Proof.

Let us consider the caterpillar tree of *n* leaf nodes. The reference clade, Λ_*R*_ has *c* number of elements. The elements of the reference clade are attached with the nodes such that the sum of the levels of those elements is minimum. For discussing here, we consider the d-node for Λ_*R*_, *dη* (Λ_*R*_), as *dη*. Let the level of the d-node (*dη*) is *p*. From Lemma 10, it can be observed that the *p* has two classes of values.

**For 0 ≤ p < c,**

- Number of leaves under *d_η_* = *n* − *p*
- Number of ingroup species placed under *d_η_* = *c* − *p*
- Number of ingroup species placed outside *d_η_* = *p*
- Number of outgroup species placed under *d_η_* = *n* − *c*

So, the Total Transfer In and Total Transfer Out are denoted as *TTL*_*in*_(*d_η_*, Λ_*R*_) and *TTL*_*out*_(*d_η_*, Λ_*R*_). Hence,

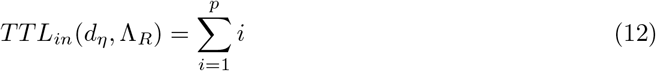

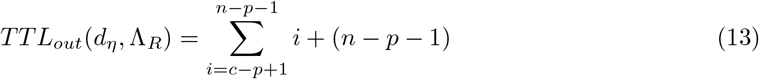

The deformation for 0 < *p* < *c* is denoted as *d*_<*c*_(*d_η_*, Λ_*R*_). Hence,

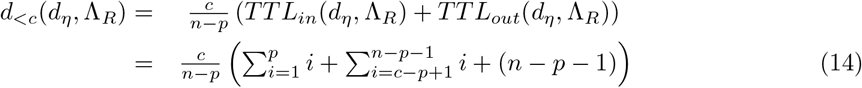

Now, for getting the value of *p* for what *d*_<*c*_(*d_η_*, Λ_*R*_) obtains the maximum value, we consider 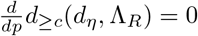. By solving this equation we get,

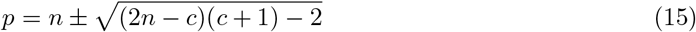

As, the maximum level of a tree of *n* leaf nodes can be (*n* − 2), so we consider,

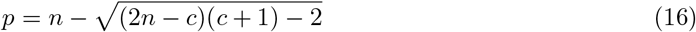

**For c** ≤ **p** < **n**,

- Number of leaves under *d*_*η*_ = *n* − *p*
- Number of ingroup species placed under *d*_*η*_ = 0
- Number of ingroup species placed outside *d*_*η*_ = *p*
- Number of outgroup species placed under *d*_*η*_ = *n* − *p*

So, the Total Transfer In and Total Transfer Out are denoted as *TTL*_*in*_(*d*_*η*_, Λ_*R*_) and *TTL*_*out*_(*d*_η_, Λ_*R*_). Hence,

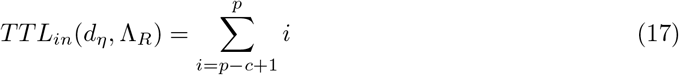

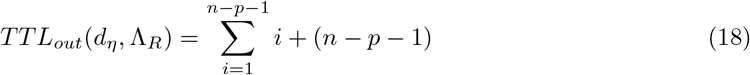

The deformation for *c* ≤ *p* < *n* is denoted as *d*_≥*c*_(*d*_η_, Λ_*R*_). Hence,

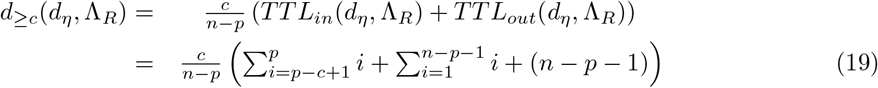

Now, for getting the value of *p* for what *d*_≥*c*_(*d*_η_, Λ_*R*_) obtains the maximum value, we consider 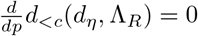. By solving this equation we get,

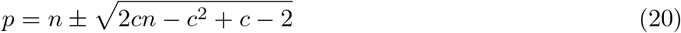

As the maximum level of a tree of *n* leaf nodes can be (*n* − 2), so we consider,

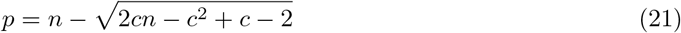

As, the level of a tree is a positive integer, hence we consider the level of the d-node as the nearest integer of *p*.

#### Corollary 1.1.

*Among all of the trees of *n* number of leaf nodes, the maximum value of the Clade Deformation for the reference clade, Λ_*R*_, of size c is denoted as Dc_max_(Λ_R_), then,*

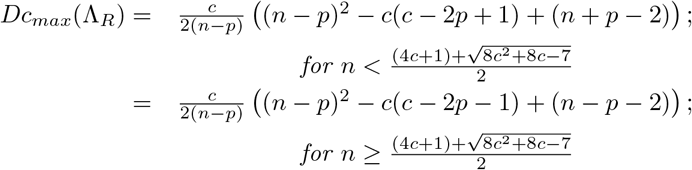

*where*,

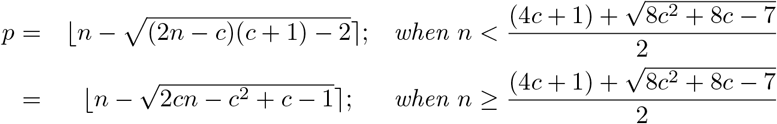

*where*, 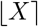 *denotes the nearest integer value of X*.

#### Proof.

Using Lemma 10 and Theorem 1 this Corollary can be easily proved.

## 5 Experimental Results and Discussion

In this section we first experimentally validate the maximum value of the Deformity Index computed in the previous section. Here we also demonstrate the distribution of the Deformity Index by computing it using a large number of random tree.

### 5.1 Yule and uniform model

We generate a large number of random trees under two different models: the Yule model [18] and the uniform model [44]. These models use two different techniques to generate random trees [29]. In the Yule model, the node is added only at the pendant edges of the base tree with an equal probability, while in the uniform model, the node is added at both internal and pendant edges of the base tree with an equal probability.

### 5.2 Simulated dataset

In our experiments we generate 100 random sets of reference clades for a given set of species. The clades of a random set together are free from the conflict relations among the species. For each set of reference clades we generate 100 trees of the same set of species randomly by considering both the Yule model and the uniform model. That way we generate 10,000 random sample points.

### 5.3 Maximum value of Clade Deformation

First we are validating Lemma 8 and 10 experimentally. Therefore, we perform an experiment where we generate a caterpillar tree of *n* leaf nodes and select random reference clades of size *k*. We computed the Clade Deformation for each random clade and considered the maximum value among them. For discussing here, we denote the maximum value of Clade Deformation for a particular *n* and *k* as *MaxC_O_*. The level of the d-node for which the maximum of the Clade Deformation occurred is considered as *MaxLυ_O_*. For the same caterpillar tree of *n* leaves, we consider *k* species having the minimum levels as the reference clade and compute the Clade Deformation and the level of the d-node. For discussing here, we denote the Clade Deformation and the level of the d-node for a particular *n* and *k* as *MaxCE* and *MaxLυ_E_*, respectively. It can be noticed that for every *n* and *k*, *MaxC_E_* > *MaxC_O_* and *MaxLυ_E_* < *MaxLυ_O_*. So it is consistent with the facts stated in Lemma 8 and 10 for all possible values of *n* and *k*. The difference between *MaxCE* and *MaxC_O_* is denoted as 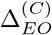 and the difference between *MaxLυ_E_* and *MaxLυ_O_* is denoted as 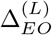. The heat maps in Fig. 6(a) and (b) show the values of 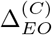 and 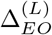 for all combinations of *n* and *k*, respectively.

**Figure 6:**
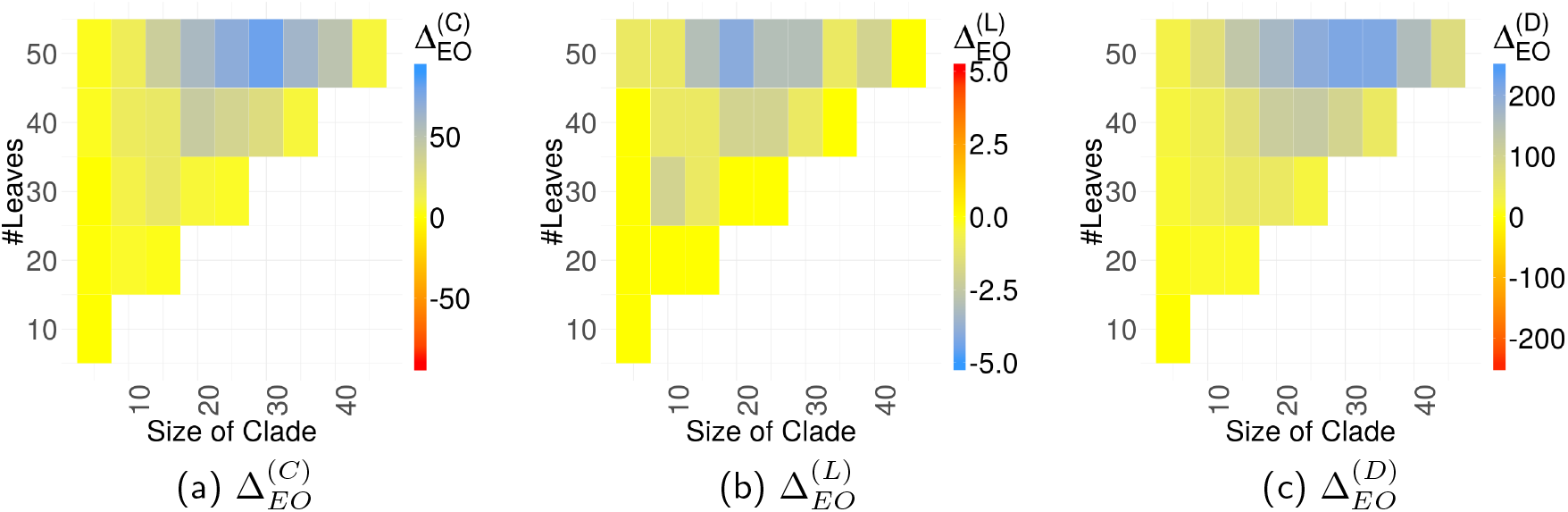
The heat maps of the differences between the expected sample and the observed sample for all combinations of *n* and *k*. (a) 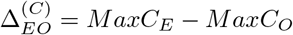 (b) 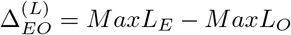 (c) 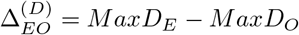

To validate Corollary 1.1, we perform an experiment where we generate random trees of a given number of leaf nodes, say *n* and also randomly select the clade of a given size, say *k*. For each value of *n* (for 10 ≤ *n* ≤ 50) and *k* (for *k* ≥ 5), we consider the maximum value of Clade Deformation among the random trees. For discussing here, we denote the maximum value of Clade Deformation for a particular *n* and *k* as *MaxD_O_*. We also derive the Clade Deformation for the tree of same configuration (*n* leaves and *k* size of clade) and the topology mentioned in Lemma 10. For discussing here, we denote it as 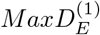. We also compute the maximum Clade Deformation from Corollary 1.1. Here, we denote it as 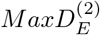. It is observed that for every *n* and *k*, 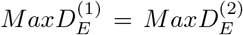, say *MaxD_E_*. So it is corroborating the fact that for the specific tree topology mentioned in Lemma 10, Corollary 1.1 is true for all possible values of *n* and *k*. It is also noticed that for all values of *n* (10 ≤ *n* ≤ 50) and *k* (*k* ≥ 5) *MaxD_O_* ≤ *MaxD_E_*. Hence, experimental results corroborate with the fact that the maximum value of Clade Deformation computed in Corollary 1.1 is satisfied for all *n* and *k*. The difference between *MaxD_E_* and *MaxD_O_* is denoted by 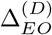. For all combinations of *n* and *k*, 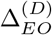 is represented by the heat map showing in Fig. 6(c).

### 5.4 Distribution of Clade Deformation

For a reference clade Λ at node *η*, the deformation (*d*(*η*, Λ)) depends on the Total Transfer In (*TTL*_*in*_(*η,* Λ)) and Total Transfer Out (*TTL*_*out*_(*η*, Λ)). From Equations 5 and 6 we find that the Transfer In and Transfer Out depend on the the level of the internal vertices. The distribution of the level of the internal vertices follows geometric distribution [51]. Since, the Total Transfer In and Total Transfer Out are linearly related with the levels of the internal vertices (refer to Equations 7 and 8), hence the Total Transfer In, Total Transfer Out, and deformation follow the geometric distribution. The deformation is finally normalized by the ratio of the size of the reference clade and the number of leaf nodes under a clade of the target tree. The distribution of the number of the leaves under a clade of a tree follows a *Beta*(1, *h*(*η*) − 1) distribution [27]. However, it is challenging to model the cumulative distribution of the Clade Deformation and remains as an open problem at this stage.

#### 5.4.1 Test for Goodness of Fit

The goodness-of-fit test is used to determine whether the observed sample distribution of a given phenomena is significantly different from the expected probability distribution. We have used two widely used techniques, Chi-Square test [36] and Kolmogorov-Smirnov (K-S) test [28], to examine whether the distribution of the Deformity Index follows a normal distribution.

In our test procedure we consider the null hypothesis as the distribution of Clade Deformation follows normal distribution. It is observed in Clade Deformation that the Chi-Square test accept the null hypothesis (with a significant level of more than 95%), but the Kolmogorov-Smirnov reject the null hypothesis.

We compute the distributions of the Deformity Index for the large number of such random trees (10,000). Fig. 7 represents the histogram of Deformity Index for the random trees generated with the 100 leaves under both the Yule model and the uniform model. It is observed that the shape of the histogram of Deformity Index for both of the two models is similar to the normal distribution. We may consider that the distribution of the Clade Deformation is close to the normal distribution, but does not follow a normal distribution.

**Figure 7:**
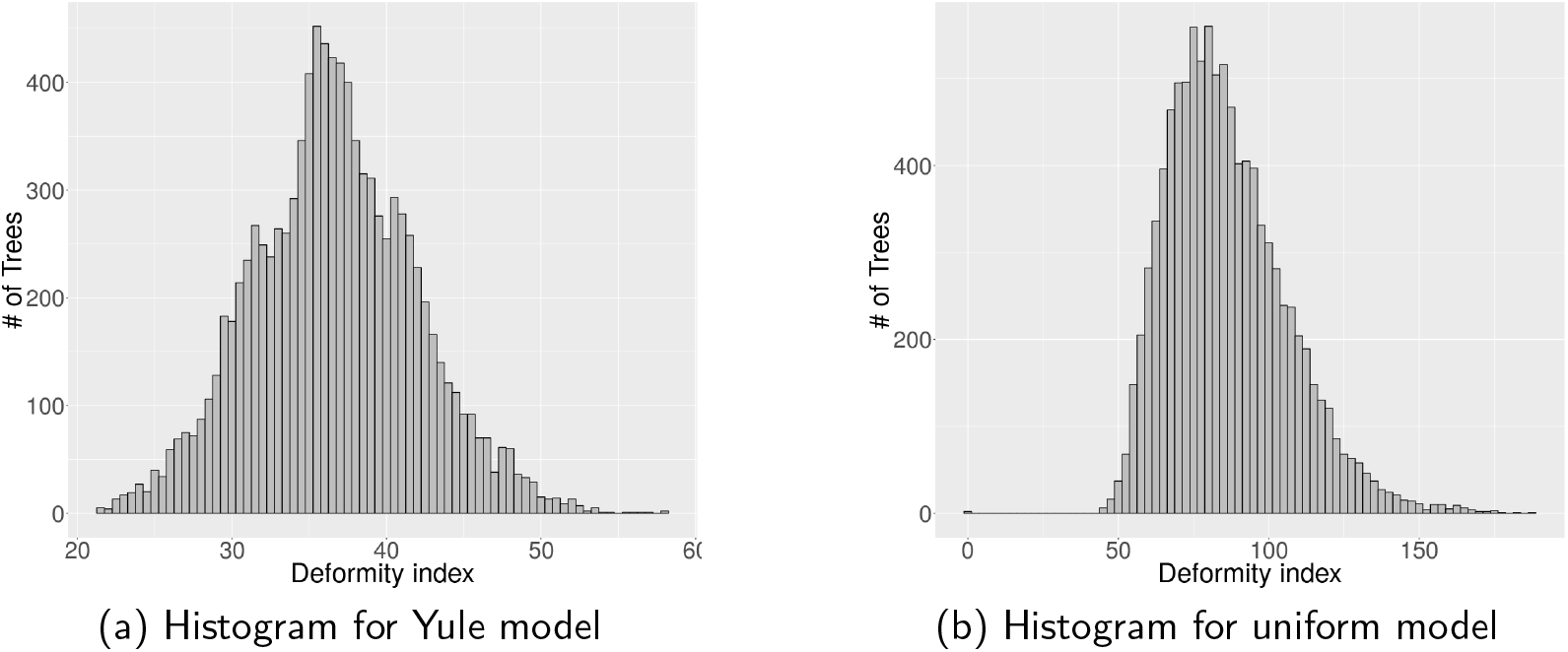
Histogram of the Deformity index for 10,000 random trees generated with 100 leaves under (a) Yule model, and (b) uniform model.

We compute the distributions of the Deformity Index for the large number of such random trees (10,000) with different number of leaves. In Fig. 8, we present the distributions of the Deformity Index for both of the Yule model and the uniform model. We can see that the mean of Deformity Index in the uniform model is larger and grow faster than that of in the Yule model. It can be also observed that the intervals of the Deformity Index in the uniform model is greater than that of in the Yule model. This phenomena represents that the random trees generated under the uniform model are more dissimilar than the random trees generated under the Yule model.

**Figure 8:**
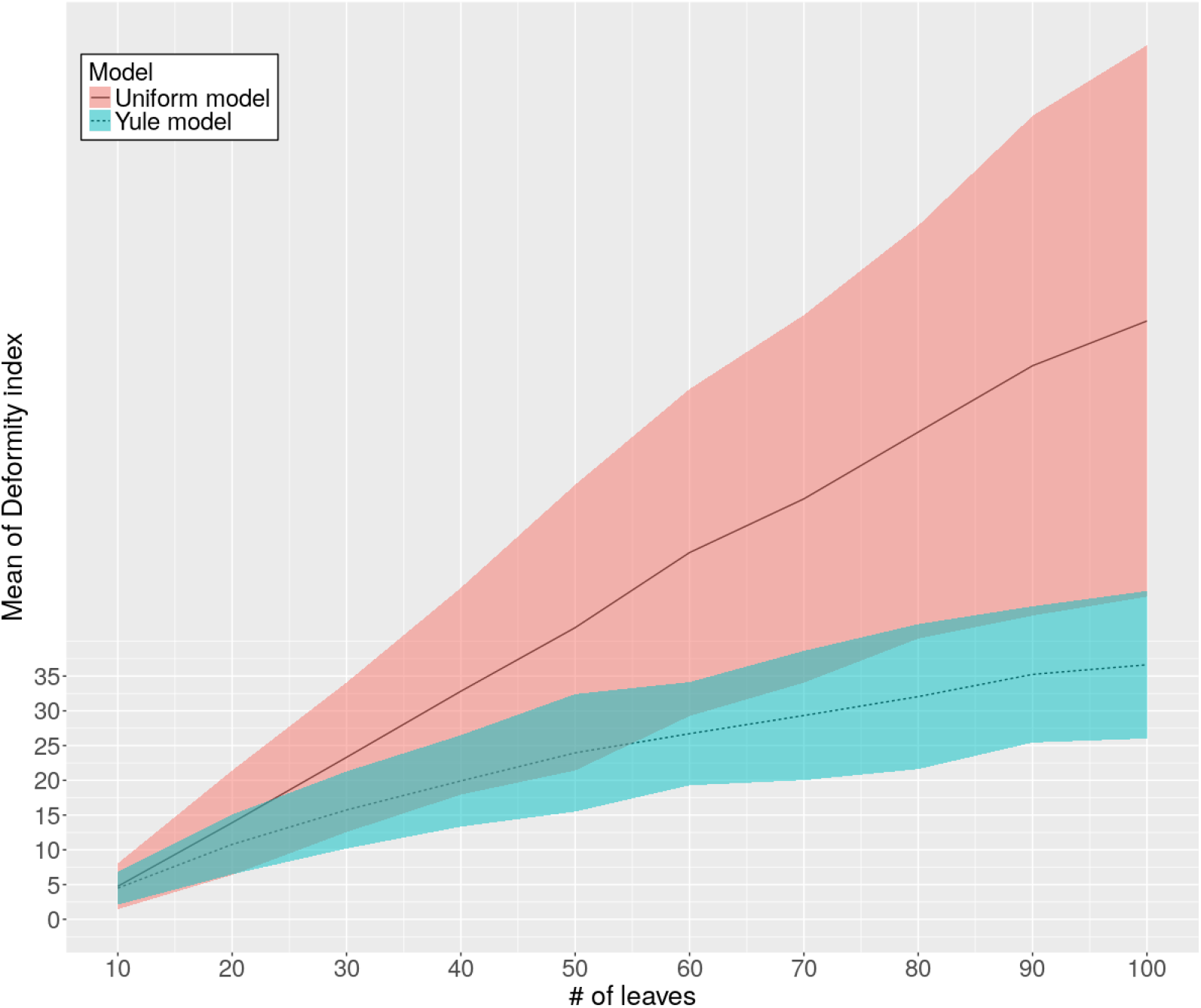
Consider the monophyletic clade has ten leaves. The distribution of the deformity index for 10,000 random trees generated under both the Yule and the uniform model with different number of leaves is shown in this figure. The solid and dashed lines show the changes of the mean of the Deformity Index w.r.t. the changes of the number of leaves for the uniform model and the Yule model, respectively. The red and blue ribbons show the double standard deviation intervals (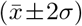) of the mean for the uniform model and the Yule model, respectively.

### 5.5 Correlation coefficients

We compute Pearson correlation coefficients [52] between Deformity Index and RF, MS, NS, and TT. The Pearson correlation coefficient (PCC) is the measure of a linear association between two sets of the same size. The value of PCC lies between −1 and +1. The positive and negative value of PCC denotes the positive and negative correlation between two variables, respectively. If value of PCC is zero, then it denotes an absence of any relationship between two variables.

The PCCs between Deformity Index and RF, MS, NS, and TT are shown in Table 1. It can be observed in Table 1 that the PCC values are very high for all the cases.

**Table 1:**
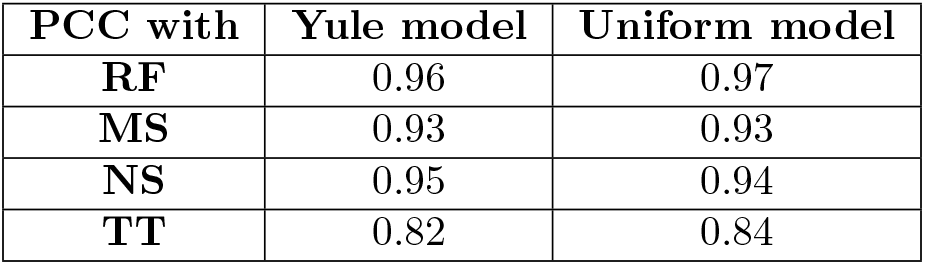
Pearson correlation coefficients between Deformity Index and RF, MS, NS, and TT for the random trees with different number of leaves generated under both the Yule model and the uniform model.

Though the proposed method is highly correlated to the conventional methods, still it can deal with the cases where the reference tree and the target tree contain different sets of species and the cases where the complete knowledge of the reference tree is not known to us. Sec. 6 clearly explains the particular scenarios where our proposed method has outperformed the existing conventional methods.

### 5.6 Complexity analysis

The deformation is the sum of *TTL*_*in*_ and *TTL*_*out*_. Its computational complexity depends on the number of species placed under the wrong clade. Considering a reference clade with *c* leaves and a tree with *n* leaves, the range of the total number of both ingroup and outgroup species placed under wrong clades is [0*, n*]. Hence, the average time complexity of computing the deformation of a clade is 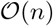. A tree with *n* leaves has the maximum of (*n* − 1) internal nodes (for binary trees). So for each reference clade, the computation of the Clade Deformation is performed for (*n* − 1) times.

Hence, the time complexity of computing the Clade Deformation of a reference clade is 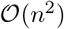. If we have *R* number of reference clades, then the time complexity of computing the Deformity Index of the tree is 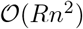.

## 6 Applications

Different studies involve in determining the phylogenetic trees of different sets of species. Hence, there are many observations on the phylogenetic relationships of the selected species. So it is very difficult to get a complete reference tree for the selected species. Reconstruction of the reference phylogenetic tree by accumulating those fragments may not be reliable. Our proposed method is more suitably applicable for those cases where the partial information of the reference tree are present. This is to be mentioned that if the complete information of the reference tree is provided to this method, then the Deformity Index will act like other conventional tree comparison techniques. There are various methods of constructing the phylogenetic tree. These methods are broadly categorized into two different categories, (i) alignment based method and (ii) alignment free method. We have examined our proposed method on the trees derived from different methods. In this section we consider two datasets of fishes and mammals.

### 6.1 Gadiformes

We choose 19 species from eight different taxonomy groups of subfamily rank of order Gadiformes and one outgroup species from order *Clupeiformes*. The details of the species are given in **Supplementary Table. 1**. We derive trees from both alignment based methods and alignment free methods.

To demonstrate the power of the Deformity Index, we consider a tree as the reference tree (shown in Fig. 9(a)) and also consider a phylogenetic tree of Gadiformes derived by maximum parsimony based method (shown in Fig. 9(b)). Both the trees have different set of species. Though these trees have visible differences, all the conventional methods of comparing those trees provide zero as the comparing score. To compute the Deformity Index, we provide all possible clades of the reference tree. The Deformity Index shows non-zero value implying that the target tree has some differences based on the reference clades.

**Figure 9:**
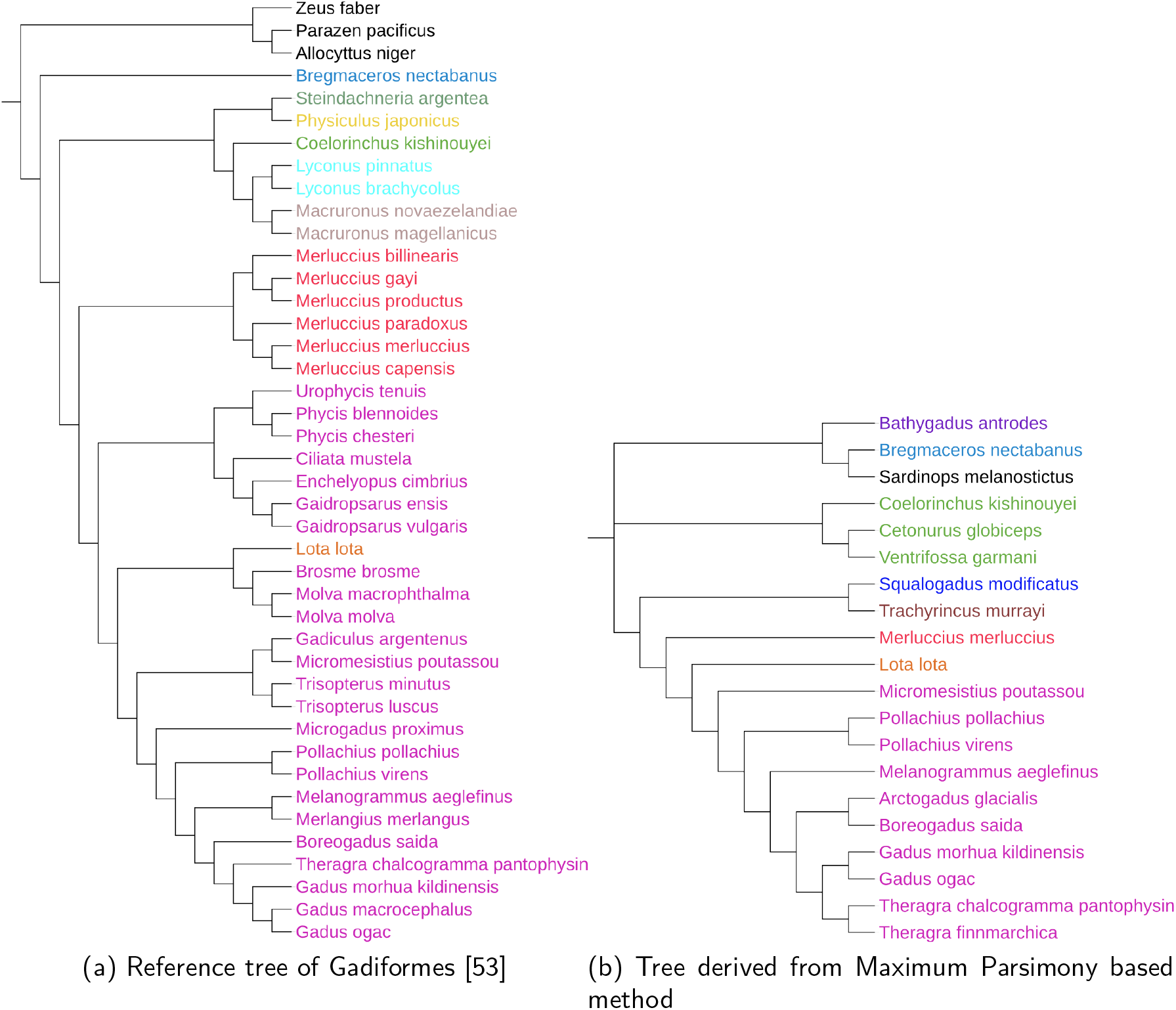
Phylogenetic tree of Gadiformes. There are different subfamilies, such as, Gadinae (pink), Lotinae (orange), Merlucciidae (red), Trachyrincinae (brown), Macrouroidae (blue), Bathygadinae (purple), Macrourinae (green), Bregmacerotidae (light blue), Lyconidae (cyan), Macruronidae (grey), Steindachneridae (light green), and Moridae (yellow). The outgroups are colored by black.

In the previous case, the reference tree gives less knowledge of relationships among the species as it has different set of species than that of the target tree. The conventional methods behave inappropriately in computing the score. This is the limitation of the conventional measures. From that point, Deformity Index takes account of the knowledge of the relationships among the species from other hypotheses and computes a score based on the cumulative knowledge. Here, we present the procedure to accumulate the knowledge of the clades from different hypotheses and use them to compute the Deformity Index.

Since long back [39] the phylogenetic relationships among the species of Gadiformes are being studied. But still there are many conflicts among various hypotheses. Hence, the true reference tree of the Gadiformes is still unknown to us. Here we consider the most accepted hypotheses of the phylogenetic relationships of these families for scoring the derived trees from different methods. The brief descriptions of the most accepted hypotheses are following. The details of these hypotheses are described in **Supplementary Sec. 1**.

***HG1*** The Gadinae subfamily is monophyletic [55, 53, 41, 45].
***HG2*** Subfamily Lotinae is the sister group of subfamily Gadinae. So, they form the monophyletic clade [33, 11, 9].
***HG3*** Bregmacerotidae belongs to the sister group of either higher “gadoids” or higher “gadoids” excluding “macruronids” [34, 41, 45]. Most of the recent studies also accept the monophyletic relation of Merlucciidae with the Gadinae and Lotinae [53, 19, 10, 45, 55]. Hence, Bregmacerotidae and Merlucciidae both form the monophyletic clade with Gadinae and Lotinae.
***HG4*** Almost all the studies agreed that all the species of Macrourinae form monophyletic clade [14].
***HG5*** The Macrouroinae and Trachyrincinae form monophyletic clade [23].
***HG6*** The Macrourinae, Macrouroinae, Trachyrincinae, and Bathygadinae form monophyletic clade [23].

We derive trees from both alignment based methods and alignment free methods. The results are given in **Supplementary Sec. 2 (Fig. 2)**. According to these hypotheses we construct the list of reference clades. Considering these reference clades we compute different quality metrics such as, Deformity Index, RF, MS, NS, and TT of the trees derived from different methods. We also judge visually the derived trees as *“complete”*, *“partial”*, and *“marginal”*, based on the reference clades.

- The *“complete”* denotes that the tree agrees the corresponding hypothesis completely.
- The *“partial”* denotes that the tree misses a few relations based on the corresponding hypothesis.
- The *“marginal”* denotes that very few relations of the hypothesis are held in the tree.

It is to be indicated that when the visual judgment is *“complete”*, then the comparing score should be zero as the tree completely supports the hypothesis while for the other cases the comparing scores should be a positive value.

For example, the tree derived from Euclidean method (refer to Fig. 10) has the monophyletic clade of the Gadinae subfamily, hence it comprises the hypothesis **HG1**completely. The Deformity Index shows zero score for that case. But all the other scores show the non-zero values because the reference clade contains the multifurcation which is not present in the derived tree (explained in Subsection 2.1). Again, it can be observed in the same tree (refer to Fig. 10) that the members of the subfamily Macrourinae form a paraphyletic clade with the other members of Gadiformes. The hypothesis **HG4** stated that the subfamily Macrourinae is a monophyletic. Hence, a high comparative score is expected. But it is observed that only the Deformity Index shows a non-zero score. All the other scores show the zero value because these methods prune all the species which do not belong to the subfamily Macrourinae (explained in Subsection 2.2). It is observed that for all of the cases values of the Deformity Index are more associated with the visual inspections than the other measures (Fig. 11). The details of this observation are given in **Supplementary Table 3**. It can be observed that the alignment-free method *D*^*^_2_ mostly corroborates to the hypotheses. The Deformity Index of the derived tree from *D*^*^_2_ shows the lowest score among all those derived trees (Table 2).

**Table 2:** Deformity Indices of the derived trees from different methods.

| Methods | Deformity Scores |  |
| --- | --- | --- |
|  | Gadiformes | Mammals |
| UPGMA [48] | 4.24 | 5.58 |
| Neighbour Joining [43] | 4.06 | 1.60 |
| Maximum Likelihood [59] | 4.06 | 1.60 |
| Maximum Parsimony [59] | 3.90 | 15.92 |
| Canberra [26] | 3.54 | 4.71 |
| Chebyshev [26] | 15.03 | 37.31 |
| Chi Square [26] | 17.84 | 21.90 |
| Co-Phylog [60] | 4.24 | 4.84 |
| Cosine [26] | 4.06 | 10.33 |
| CV Tree [38, 63] | 4.81 | **0.13** |
| $D_2^*$ [40, 57] | **3.34** | 11.01 |
| Euclidean distance [26] | 10.26 | 19.38 |
| FFP [46, 47] | 24.01 | 26.50 |
| Fast Vector [25] | 11.64 | 5.29 |

**Figure 10:**
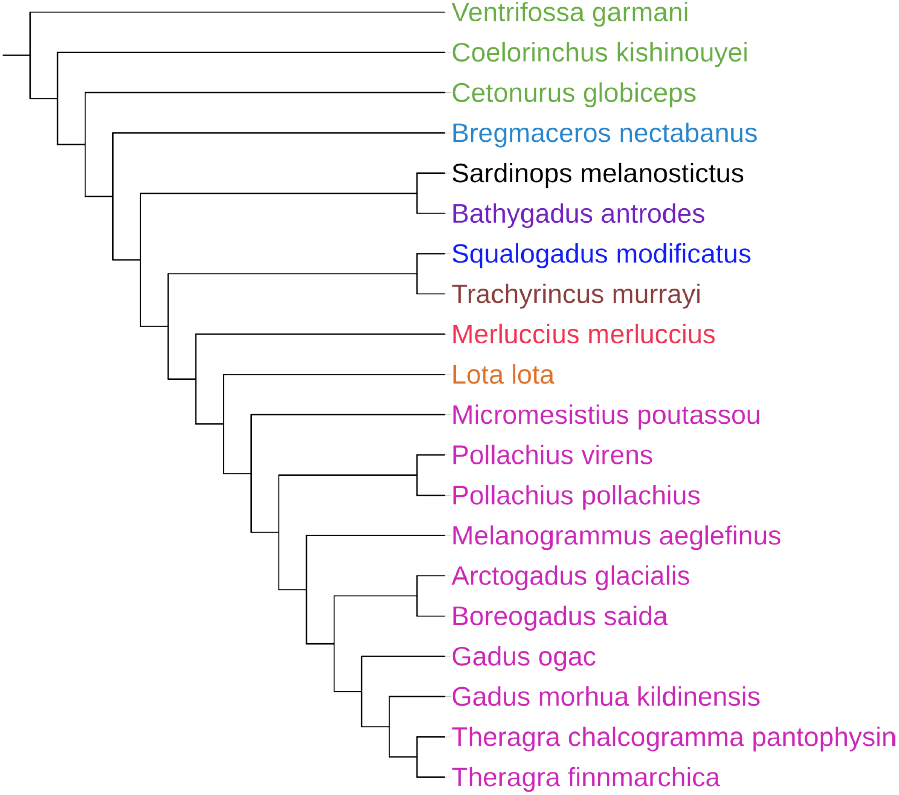
Phylogenetic tree of Gadiformes derived by Euclidean method. There are different subfamilies, such as, Gadinae (pink), Lotinae (orange), Merlucciidae (red), Trachyrincinae (brown), Macrouroidae (blue), Bathygadinae (purple), Macrourinae (green), Bregmacerotidae (light blue). The outgroup are colored by black.

**Figure 11:**
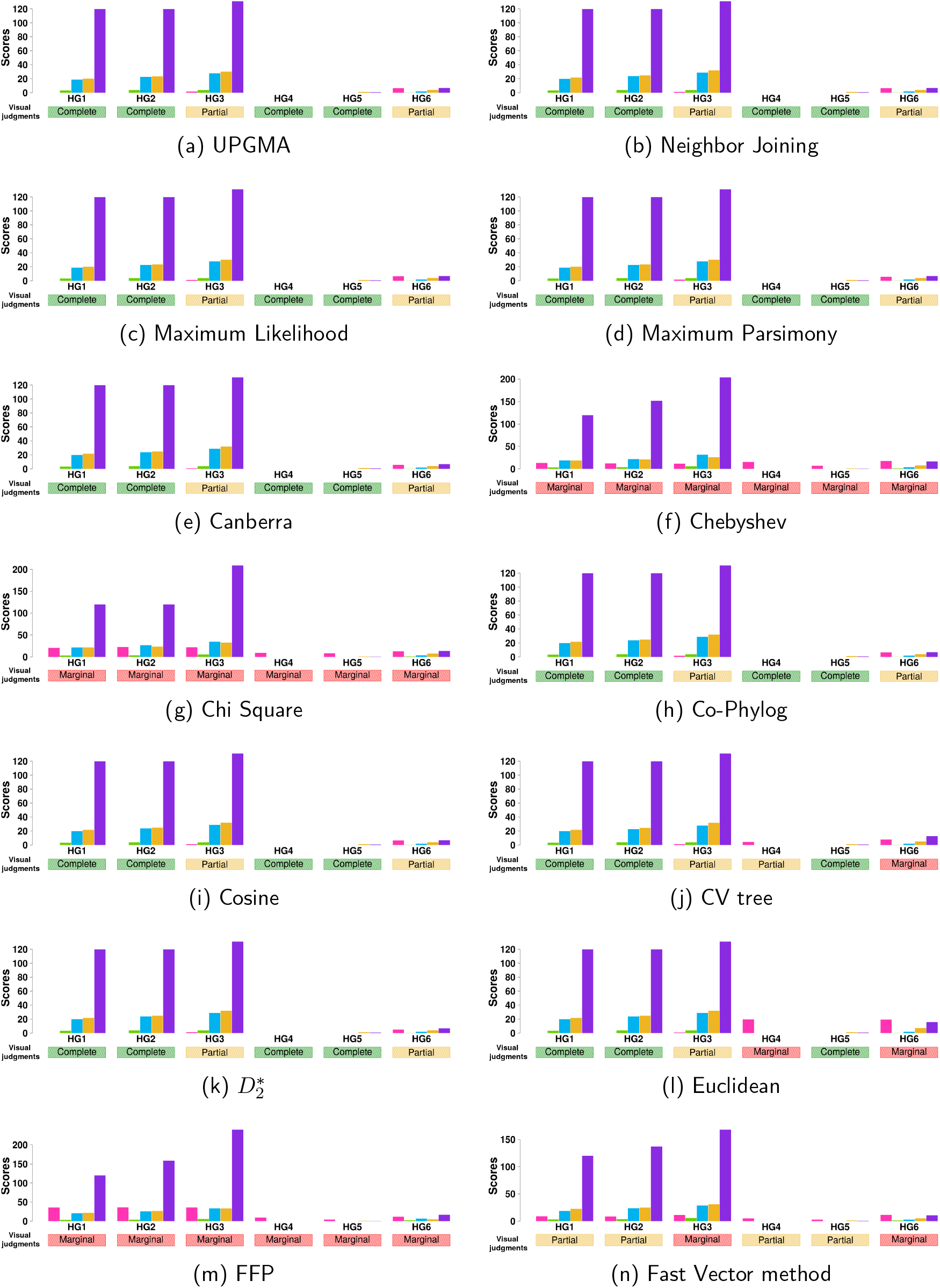
Comparative scores of the trees derived from different methods for different hypotheses. For Gadiformes, the scores shown here are Deformity Indices (pink), RF (green), MS (blue), NS (yellow), and TT (purple), respectively. The reference clades or the hypotheses are described as **HG1 - HG6**. The non-existence of a bar denotes the zero value of the corresponding score. It is to be mentioned that for the visual judgment *“complete”*, the scores should be zero while for the *“partial”* and *“marginal”* cases the scores should be nonzero positive values. It is shown that the deformity index (pink) consistently satisfies this property, while the other scoring methods underperform in many cases in this respect.

### 6.2 Mammals

We consider 40 mammals from seven orders of class Mammalia. The detail of the species are given in the **Supplementary Table 2**. The phylogeny of mammals are highly studied over the years [30, 31, 50, 56, 22, 17, 32, 37]. But there are many different hypotheses available in the literature of the phylogeny of mammals. For computing the Deformity Index of the derived trees, we consider some of the widely accepted hypotheses. The hypotheses are described in detail in the **Supplementary Sec. 3**. A few of these widely accepted hypotheses is listed below.

***HM1*** The order Primates forms a monophyletic clade [25, 30, 31, 56, 22, 17]
***HM2*** Human is the sibling of chimpanzee and gorilla is the sister of (human + chimpanzee) [17, 37].
***HM3*** The order Carnivora form the monophyletic clade [25, 56, 50, 30, 31].
***HM4*** Most of the studies supported the order Rodentia as the sister of Lagomorpha [30, 31, 56, 50].
***HM5*** Most of studies agreed that Rodentia and Lagomorpha are sisters of Primates [30, 31, 50].
***HM6*** Most of the studies supported that the order Perissodactyla is the sister group of order Carnivora [30, 31, 50].

Both the alignment based and alignment free methods are implemented on the mammals dataset. It is also to be mentioned that considering the same dataset we found some conflicts between the result provided in [25] and the tree obtained by applying the Fast Vector method (please refer to Fig. 12).

**Figure 12:**
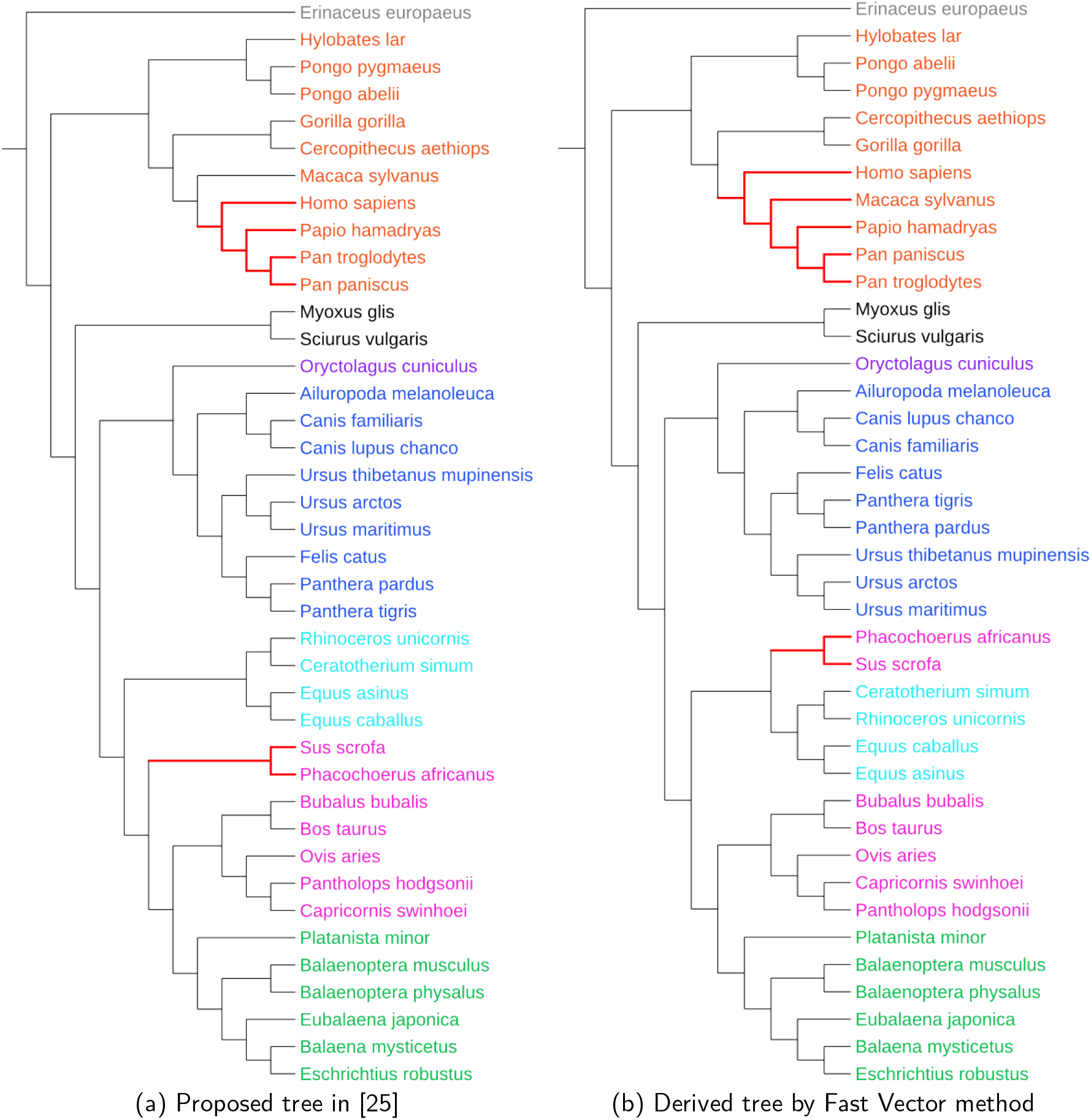
Both the trees are constructed using 40 species of mammals. They are grouped as seven orders. The Cetartiodactyla again divided into Artiodactyla and Cetacea. The order are Primates (red), Cetacea (green), Artiodactyla (pink), Perissodactyla (light green), Rodentia (black), Lagomorpha (dark red), Carnivore (blue), and Erinaceomorpha (grey). (a) Tree proposed in [25] which was derived from their proposed method Fast Vector. (b) The generated tree by applying the Fast Vector on the same dataset. The discrepancies between these two trees are shown in Red colored clade.

The details of the derived tree from each of the methods are provided in **Supplementary Sec. 4 (Fig. 3)**. Based on these studies the reference clades can be prepared to compute the Deformity Index of the trees derived from different methods. Similar to the Gadiformes we also consider these reference clades and compute the Deformity Index, RF, MS, NS, and TT measures of the trees derived from different methods. We judge visually the derived trees as “complete”, “partial”, and “marginal”, based on the reference clades, as we have done for Gaidformes. The tree derived from Fast Vector method (refer to Fig. 12(b)) contains the monophyletic clade of Carnivora, so it supports the hypothesis **HM3**. Hence, a zero comparative score is expected. But it is noticed that except the Deformity Index all the other scoring methods show the non-zero values because the reference clade depicts the multifurcation relationships which are not present in the derived tree (explained in Subsection 2.1). In another case, the same tree (refer to Fig. 12(b)) has partial support for both the hypotheses **HM1** and **HM4**. Therefore, a non-zero scores are expected for both of the cases. But it is observed that only the Deformity Index shows the non-zero values for both of the cases. The other methods prune those species which do not belong to the corresponding order and shows the zero scores (explain in Subsection 2.2). It is also observed that for all of the cases values of the Deformity Index are more associated with the visual inspections than the other measures (Fig. 13). The detail of this observation is given in **Supplementary Table 4**. It is observed that based on the hypotheses discussed here, the CV tree method gives better result than all of the other methods. The Deformity Index of the tree derived from the CV tree method shows the lowest score than that of the other methods (Table 2).

## 7 Conclusion

In this paper, we propose a new semi-reference method to measure the quality of a tree using the biological knowledge of the clades. The Deformity Index of the tree gives an idea about the correctness of the clades within the tree. As this method only depends on the biological knowledge of the clades, so the Deformity Index can easily adapt with the present knowledge in biology and provides the quality metric in that context. At the same time, Deformity Index can also adapt itself in versatile scenarios where the other conventional methods of comparing trees lack to provide a meaningful comparison scores (described in Fig. 1). The complexity of computing the Deformity Index is 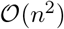, which is also comparable with the other conventional methods. We have studied different properties of the Deformity Index, such as, minimum and maximum values. We inspect the distributions of different modules of the Deformity Index. We also perform various statistical tests, such as, Chi-Square test and Kolmogorov-Smirnov test for the goodness of fit to understand the distribution of Deformity Index. We have given extremal results as well as experimental results for different biological models to characterize the proposed method. Considering two datasets of fishes and mammals, we apply this measure to score the biological trees generated by different state of the art methods. It is observed that higher their degree of adherence to the widely accepted hypotheses about various phylogeny, lower is the score of their Deformity Index.

**Figure 13:**
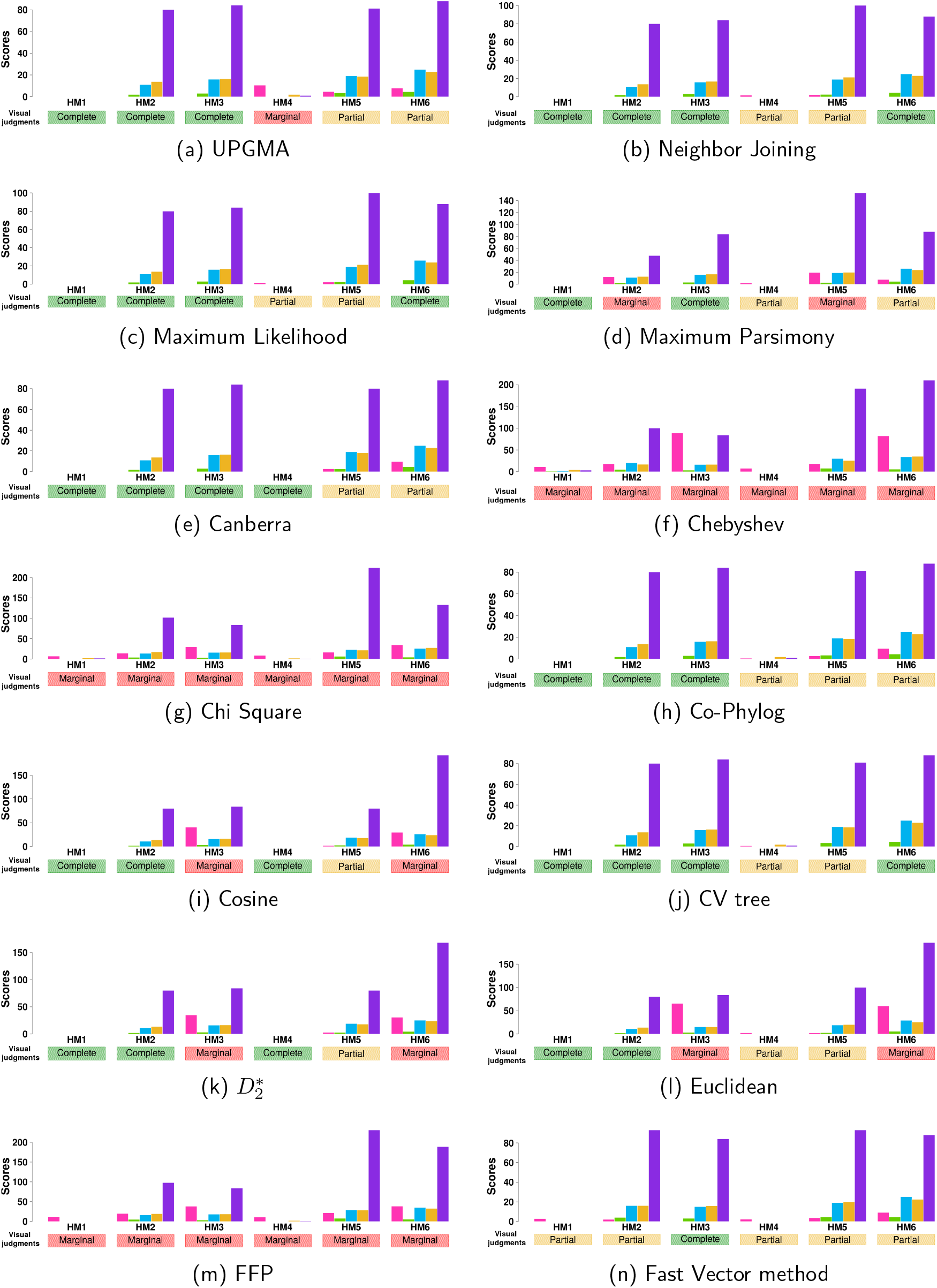
Comparative scores of the trees derived from different methods for different hypotheses. For mammals, the scores shown here are Deformity Indices (pink), RF (green), MS (blue), NS (yellow), and TT (purple), respectively. The reference clades or the hypotheses are described as **HM1 - HM6**. The non-existence of a bar denotes the zero value of the corresponding score. It is to be mentioned that for the visual judgment *“complete”*, the scores should be zero while for the *“partial”* and *“marginal”* cases the scores should be nonzero positive values. It is shown that the deformity index (pink) consistently satisfies this property, while the other scoring methods underperform in many cases in this respect.

## Appendix

### Proof.

The level of a node is the number of edges present between root and that corresponding node of the tree. Multifurcation at a node reflect the co-occurrence of multiple nodes at the same node. This phenomena reduces the number of edges at the path. Hence, any occurrence of multifurcation on the path from root to a node reduce the level of that node. So, it can be said that the level of a node of a tree will be maximum if and only if all the ancestors of the node are bifurcated.

### Lemma 7.

*For a given number of leaf nodes, the highest value of the sum of the levels of all the leaves of a tree achieves when the tree is a caterpillar tree*.

### Proof.

From Lemma 6 it is understood that the maximum value of the level of a node occurs when the tree is a binary tree. Hence,

***Base case:*** For the number of leaf nodes of a tree, *n* = 3, the tree is a caterpillar tree. The sum of the levels of all the leaves achieves the highest value for that topology.

***Induction step:*** Let us consider a caterpillar tree with *k* ∈ Z+ number of leaf nodes which have the highest value of the sum of the levels of all its leaves. Our target is to add a new leaf node to the tree and also get the highest value of the sum of the levels of all the leaves among all the trees having (*k* + 1) number of leaf nodes.

The height of a tree is proportional to the levels of the leaf nodes of the tree. Hence, we have to attach the new node such that the addition of the new node will increase the height of the tree. This can only be possible if the attachment of the new node also form a caterpillar tree.

Thus, Lemma 7 holds for *n* = *k* + 1.

***Conclusion:*** By the principle of induction, Lemma 7 is true for all *n* ∈ Z+.

### Lemma 8.

*For a caterpillar tree, lower the sum of the levels of the elements of reference clade is equivalent to higher the distortion of the clade within the tree*.

### Proof.

Let us consider a caterpillar tree of *n* number of leaf nodes and consider a reference clade, Λ_*R*_, of size *c*.

For the ideal case, all the elements of the reference clade form a monophyletic clade. So the outgroup elements should be placed out side of the clade. For a caterpillar tree it can only be possible if all the outgroup elements are placed at the upper level of the elements of the reference clade.

Therefore, for the ideal cases, *h*(*η*_*in*_) > *h*(*η*_*out*_), ∀*η*_*in*_ ∈ Λ_*R*_, ∀*η*_*out*_ ∉ Λ_*R*_, and *h*(*n*) is the level of the node *n*. Hence, for that topology, the sum of the levels of the elements of the reference clade is maximum.

Now if we exchange the position of an element of the reference clade with an outgroup element, the sum of the levels of the ingroup elements will be reduced. More we exchange the position between ingroup and outgroup elements, more we reduce the sum of the level of ingroup elements.

So, we can conclude that for the caterpillar tree the extreme distortion of the reference clade will occur while the sum of the levels of the ingroup elements is minimum.

### Proof.

If we can prove that once the deformation starts increasing at the *i*^*th*^, ∀*i* ≥ 0 level, it never decreases its value at any of the nodes of the level > *i*, then we can draw the implication that the deformation is a convex function.

Let us consider, a reference clade Λ with *k* species of a caterpillar tree with *n* leaves. The internal nodes of levels *i*, *i* + 1, and *i* + 2 are denoted as *η*_*i*_, *η*_*i*+1_, and *η*_*i*+2_, respectively.

**At the node** *η*_*i*_, let us consider the followings,

- Number of leaf nodes under *η*_*i*_ = *l*
- Number of ingroup species placed under *η*_*i*_ = *m*
- Number of outgroup species placed under *η*_*i*_ = *n*
- *TTL*_*in*_(*η*_*i*_, Λ) = *X*_*in*_
- *TTL*_*out*_(*η*_*i*_, Λ) = *Y*_*out*_

Hence, 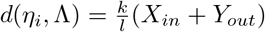

**At the node** *η*_*i*+1_, In caterpillar tree, increasing a level through the internal node removes a leaf node from the clade of previous level. There can two cases,

- **Case 1:** removed leaf is an outgroup species
- **Case 2:** removed leaf is an ingroup species

For Case:1 we can derive the followings,

- Number of leaf nodes under *η*_*i*+1_ = *l* − 1
- Number of ingroup species placed under *η*_*i*+1_ = *m*
- Number of outgroup species placed under *η*_*i*+1_ = *n* − 1
- *TTL*_*in*_(*η*_*i*+1_, Λ) = *Xin* + (*k* − *m*)
- *TTL*_*out*_(*η*_*i*+1_, Λ) = *Yout* − (*n* − 1)

For Case:2 we can derive the followings,

- Number of leaf nodes under *η*_*i*+1_ = *l* − 1
- Number of ingroup species placed under *η*_*i*+1_ = *m* − 1
- Number of outgroup species placed under *η*_*i*+1_ = *n*
- *TTL*_*in*_(*η*_*i*+1_, Λ) = *Xin* + (*k* − *m* + 1)
- *TTL*_*out*_(*η*_*i*+1_, Λ) = *Yout* − *n*

Hence, for both the cases,

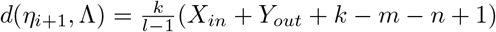

Similarly, **at the node** *η*_*i*_,

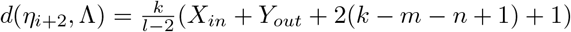

As we consider that the deformation is increasing then *d*(*η*_*i*_, Λ) < *d*(*η*_*i*+1_, Λ). Hence,

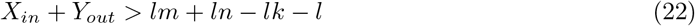

Now for the case when deformation decreases at the level *i* + 1, then *d*(*η*_*i*+1_, Λ) > *d*(*η*_*i*+2_, Λ). Therefore,

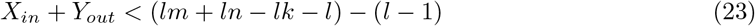

As both the Equations 22 and 23 can not be satisfied simultaneously for any value of *l* > 1, so we can claim that the deformation is a convex function for the caterpillar tree.

### Lemma 10.

*For a caterpillar tree of n number of leaves and a reference clade of size c* (1 < *c* < *n*), *the level of the d-node is p*, *where*,

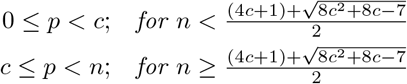

### Proof.

In Lemma 7, we have proved that for a given number of leaf nodes, the maximum level of each leaves occur when the tree is a caterpillar tree. Lemma 8 proved that the distortion of a reference clade will be maximum while the sum of the levels of all the elements of the reference clade is minimum. Hence, to prove the Lemma 10, we consider a caterpillar tree and the elements of the reference clade is placed at their minimal level.

For a caterpillar tree of topology described in Lemma 8, let the level of d-node be, *h*(*d*_*η*_ (Λ_*R*_)) = *p*. If we move through the internal nodes from root, the deformation score first monotonously decreases till d-node and reach its minimum value. Then after d-node, it again increases monotonously till the highest level (refer to Lemma 9).

Consider a caterpillar tree of leaves *n* and a reference clade, Λ_*R*_, of size *c*. Let us consider the deformation of Λ_*R*_ at node *η*_1_ of level (*c* − 1) and at node *η*_2_ of level *c* of that caterpillar tree as *d*(*η*1, Λ_*R*_) and *d*(*η*_2_, Λ_*R*_), respectively. Suppose, the d-node appears before the level *c*. Hence, *p* < *c*.

**At node** *η*_**1**_ **of level(c − 1)**,

- Number of leaves under *η*_1_ = *n* − *c* + 1
- Number of ingroup species placed under *η*_1_ = 1
- Number of ingroup species placed outside *η*_1_ = *c* − 1
- Number of outgroup species placed under *η*_1_ = *n* − *c*

So, the Total Transfer In and Total Transfer Out are denoted as *TTL*_*in*_(*η*_1_, Λ_*R*_) and *TTL*_*out*_(*η*_1_, Λ_*R*_). Hence,

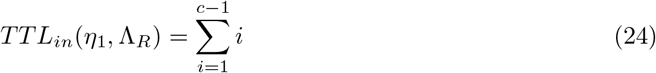

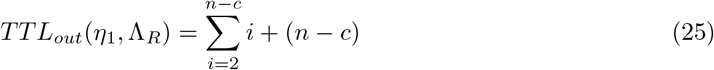

Hence,

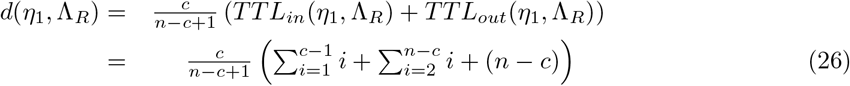

**Similarly, at node** *η*_**2**_ **of level c**,

- Number of leaves under *η*_2_ = *n* − *c*
- Number of ingroup species placed under *η*_2_ = 0
- Number of ingroup species placed outside *η*_2_ = *c*
- Number of outgroup species placed under *η*_2_ = *n* − *c*

So, the Total Transfer In and Total Transfer Out are denoted as *TTL*_*in*_(*η*_2_, Λ_*R*_) and *TTL*_*out*_(*η*_2_, Λ_*R*_). Hence,

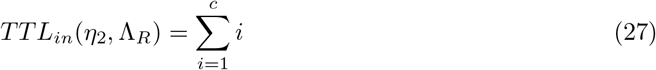

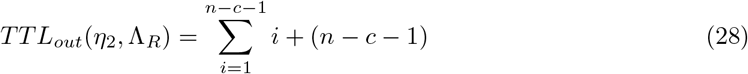

Hence,

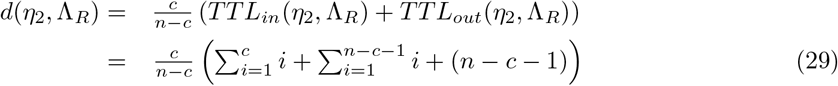

As *p* < *c*, then *d*(*η*_1_, Λ_*R*_) < *d*(*η*_2_, Λ_*R*_).

Therefore, *d*(*η*_1_, Λ_*R*_) < *d*(*η*_2_, Λ_*R*_) while,

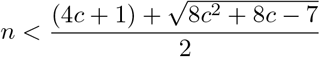

Hence, for 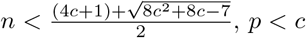

